# *Streptococcus agalactiae* and *Escherichia coli* Induce Distinct Effector γδ T Cell Responses During Neonatal Sepsis

**DOI:** 10.1101/2023.10.02.560561

**Authors:** Lila T. Witt, Kara G. Greenfield, Kathryn A. Knoop

## Abstract

Neonates born prematurely are highly vulnerable to life-threatening conditions such as bacterial sepsis. *Streptococcus agalactiae*, also known as group B *Streptococcus* (GBS) and *Escherichia coli* are frequent causative pathogens of neonatal sepsis, however, it remains unclear if distinct sepsis pathogens induce differential adaptive immune responses. In the present study, we find that γδ T cells in neonatal mice rapidly respond to single-organism GBS and *E. coli* bloodstream infections and that these pathogens induce distinct activation and cytokine production from IFN-γ and IL-17 producing γδ T cells, respectively. We also report differential reliance on γδTCR signaling to elicit effector cytokine responses during neonatal sepsis, with IL-17 production during *E. coli* infection being driven by γδTCR signaling, and IFN-γ production during GBS infection occurring independently of γδTCR signaling. Furthermore, we report that the divergent effector responses of γδ T cells during GBS and *E. coli* infections impart distinctive neuroinflammatory phenotypes on the neonatal brain. The present study reveals that the neonatal adaptive immune system differentially responds to distinct bacterial stimuli, resulting in unique neuroinflammatory phenotypes.

## Introduction

Premature neonates are acutely at risk for the development of life-threatening infections such as neonatal sepsis and meningitis.^1,2^ Neonatal susceptibility to infection is not only facilitated by environmental factors such as prolonged stay in the neonatal intensive care unit (NICU) but also by the relative immaturity of the neonatal immune system.^1,3,4^ Although the understanding of the neonatal immune system has progressed significantly over the past decade, how the neonatal adaptive immune system responds to disparate bacterial insults remains poorly understood.

Bloodstream infections preceding neonatal sepsis and meningitis can be caused by both Gram-negative and Gram-positive bacteria. *Escherichia coli (E. coli)* and *Streptococcus agalactiae* (Group B *Streptococcus*; GBS) are among the most common causative pathogens implicated in neonatal bacterial sepsis.^5^ Gram-negative bacilli such as *Klebsiella*, *Pseudomonas,* and *Escherichia coli (E. coli)* are prevalent in the gastrointestinal tract of premature neonates and are capable of translocating from the gut and causing sepsis.^3,6,7^ Increased bacterial translocation from the neonatal gut is facilitated in part by selective deficiencies in gut barrier defense mechanisms, including decreased production of protective factors such as mucous, anti-microbial peptides, and IgA.^6^

Conversely, Gram-positive neonatal sepsis is frequently caused by *Staphylococcus aureus* and *Streptococcus agalactia*e, or group B *Streptococcus* (GBS). GBS colonizes the neonate via vertical transmission during birth, often as the result of the neonate aspirating GBS-infected amniotic fluid during parturition.^8^ Although rates of GBS neonatal sepsis are declining due to improved early detection methods and prophylactic maternal antibiotic administration, the mortality rate of GBS neonatal sepsis can be as high as 10%, with 30-50% of survivors going on to experience neurological comorbidities in early childhood.^9^

Although both GBS and *E. coli* can be found as components of a healthy microbiota, they have the potential to cause severe disease in vulnerable populations, such as neonates.^10^ The increased risk of sepsis development amongst preterm neonates is further compounded by deficiencies in several innate immunological defense mechanisms. Compared to adults, neonates have reduced complement proteins in their blood, impaired neutrophil function, , and impaired secretion of pro-inflammatory cytokines by dendritic cells.^11^ Similar to the innate immune system, the adaptive immune system in neonates bears several striking deficiencies to that of adults,^11^ including a skewing of CD4+ T cells toward Th2 over Th1 differentiation, further impairing the ability of neonates to mount a proper immune response to microbial infections.^12–14^ Neonates also have deficiencies in humoral immunity, such as delayed germinal center formation and reduced antibody responses to both T cell-dependent and independent antigens.^15^

Despite their relative impairments in the conventional T and B cell compartments, neonates have a functional population of γδ T cells.^13^ γδ T cells are innate-like lymphocytes that are abundant in barrier sites and act as early immune sentinels during infection.^16,17^ In contrast to conventional CD4+ and CD8+ T cells, γδ T cells are exported from the thymus as functionally mature cells and are poised to rapidly deploy their effector functions upon the detection of microbial ligands or pro-inflammatory cytokines.^18^ As the first T cells to develop in the embryonic thymus,^19,20^ γδ T cells are critical players in the neonatal immune response during a time when CD4+ and CD8+ T cells, and B cells are still developing and maturing.^17,21,22^ Indeed, γδ T cells have been shown to play a critical role in host protection during neonatal influenza ^23^ and *Clostridium difficile* infection ^24^ underscoring their importance during early life.

In the present study, we characterize the immune responses to two major neonatal sepsis pathogens, *Streptococcus agalactiae* (Group B *Streptococcus*) and *Escherichia coli.* We report that these two pathogens induce distinct effector cytokine responses from γδ T cells in postnatal day 7 (P7) pups. We also report that these two pathogens drive distinct neuroinflammatory phenotypes in neonatal mice and that γδ T cells contribute to sepsis-induced neuroinflammation in a pathogen-specific manner. This study sheds light on how distinct sepsis pathogens drive differential γδ T cell effector responses in neonatal mice.

## Results

### *γδ T cells Respond to* E. coli *and GBS Neonatal Sepsis*

While group B *Streptococcus* (GBS) and *Escherichia coli* are frequent causative pathogens of neonatal sepsis, it is still unclear if features of these bacteria differentially drive the neonatal adaptive immune response. Previous studies have suggested that *GBS* and *E. coli* may utilize different routes of entry to cause sepsis: neonatal GBS disease is often the result of vertical transmission *in utero* or during parturition^25,26^; however, clinical data has suggested that GBS can also disseminate from the gut to cause sepsis.^27^ *E. coli* is also associated with enteric sepsis, as we and others have shown that *E. coli* can readily translocate from the neonatal intestine to cause sepsis.^3^ Therefore, we intraperitoneally infected neonatal pups (postnatal day 7; P7) with a single-organism infection of either 10^6^ CFU *Streptococcus agalactiae* (GBS) or 2×10^4^ CFU *E. coli* “Bloodstream Isolate B” (BSI-B) to directly compare the systemic response to these bacteria independent of the route of entry. Pups were sacrificed at 18 hours post-infection, a time point at which sepsis-induced weight loss begins (data not shown), and the spleen was analyzed as a readout of the systemic inflammatory response to GBS and *E. coli* infection. Both *E. coli* and GBS septicemia induced robust activation of splenic γδ T cells 18 hours post-infection, as measured by an increase in the proportion of activated (CD69+ CD62L-) γδ T cells compared to uninfected controls (Fig. 1a, b). A significant increase in the absolute number of activated γδ T cells during *E. coli* infection was also observed, along with a modest but statistically non-significant increase in the number of activated γδ T cells during GBS infection (Fig. 1c). In addition to the γδ T cell compartment, a slight increase in the activation status of conventional CD4+ and CD8+ T cells, and B cells was also observed (Fig. S1a-c). Therefore, these data show that γδ T cells in the neonate rapidly respond to both *E. coli* and GBS infection.

**Figure 1:**
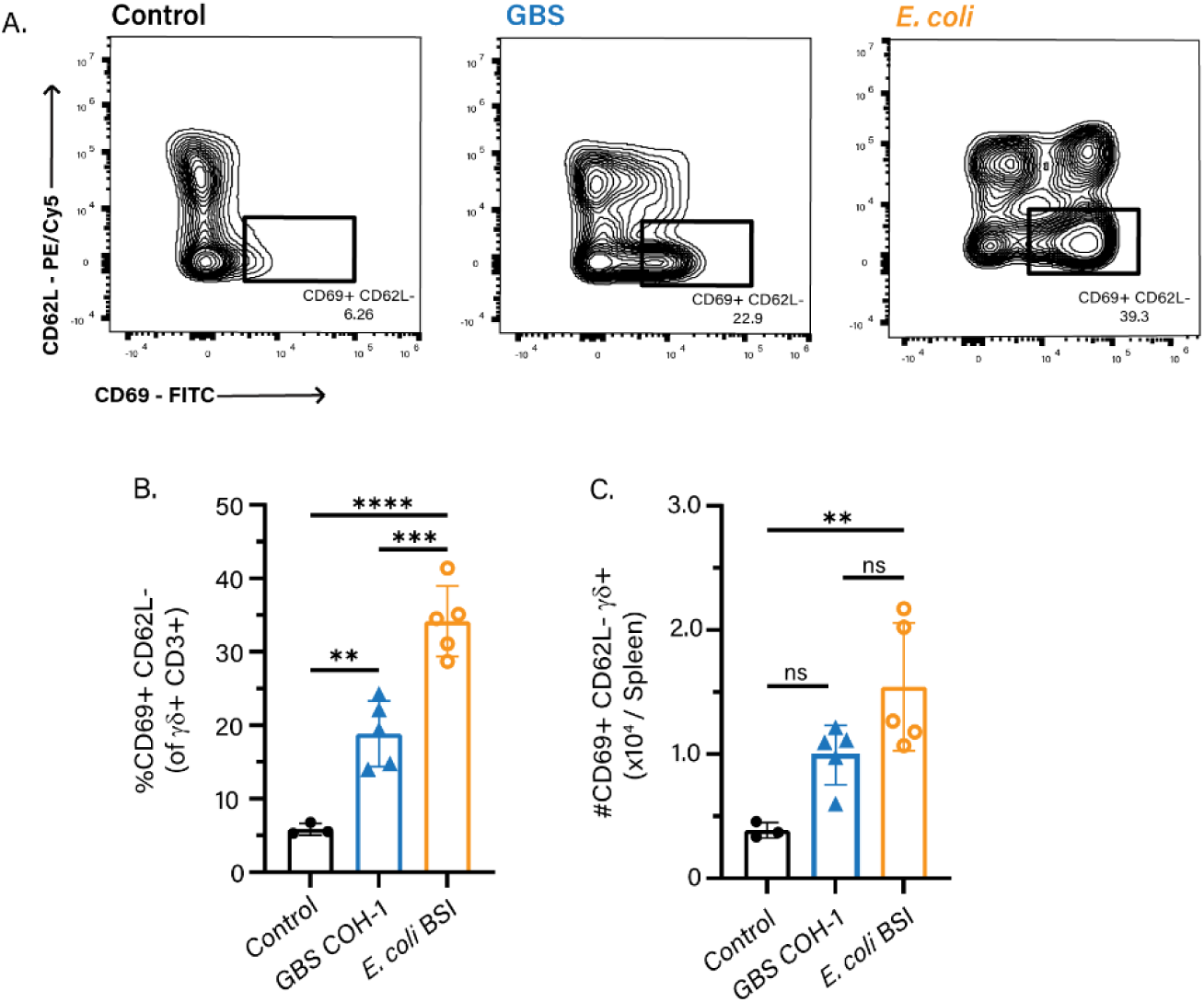
γδ T cells Respond to *E. coli* and GBS Neonatal Sepsis. BL/6 pups were infected with GBS or *E. coli* on P7, and the spleen was analyzed 18 hours post-infection. A) Gating scheme, B) frequency, and C) absolute number of CD69+, CD62L-splenic γδ+ T cells. The data shown is from four independent experiments with n>3 mice per group, where each dot represents one mouse. Controls are uninfected age-matched littermates. Statistical tests used include one-way ANOVA (B, C) with ns = p>0.05, ** = p ≤ 0.01, *** = p ≤ 0.001, **** = p ≤ 0.0001.

### E. coli and GBS Neonatal Sepsis Drive Distinct Effector Cytokine Responses from γδ T cells

We next sought to further characterize the responses of γδ T cells during GBS and *E. coli* neonatal sepsis. γδ T cells undergo developmental programming in the thymus and exist in the periphery as either IL-17 or IFN-γ producers.^18,28,29^ ^18^. We therefore asked which γδ T cell effector cytokine profiles were being elicited during neonatal GBS and *E. coli* infection. Cytokine staining of total splenic γδ T cells revealed a robust increase in IL-17 expression during *E. coli*, but not GBS infection, whereas GBS infection induced increased IFN-γ, but not IL-17, expression from γδ T cells (Fig. 2a-c). These findings were validated with serum ELISA, showing global systemic increases in IFN-γ during GBS infection and IL-17 during *E. coli* infection (Fig. 2d, e). γδ T cell effector programs can also be discerned based on the expression of surface markers, such as CCR6, restricted to IL-17 producing γδ T cells, and CD27, restricted to IFN-γ producing γδ T cells. Accordingly, during GBS infection, there was an increase in the proportion of activated CD27+ γδ T cells, but not CCR6+ γδ T cells compared to uninfected controls (Fig. 2f). Conversely, during *E. coli* infection, there was an increase in the proportion of activated CCR6+ γδ T cells, but not CD27+ γδ T cells compared to uninfected controls (Fig. 2g). These data demonstrate that GBS neonatal sepsis drives the specific activation of CD27+, IFN-γ-producing γδ T cells, whereas *E. coli* infection activates CCR6+, IL-17-producing γδ T cells, resulting in the induction of discrete effector cytokine programs.

**Figure 2:**
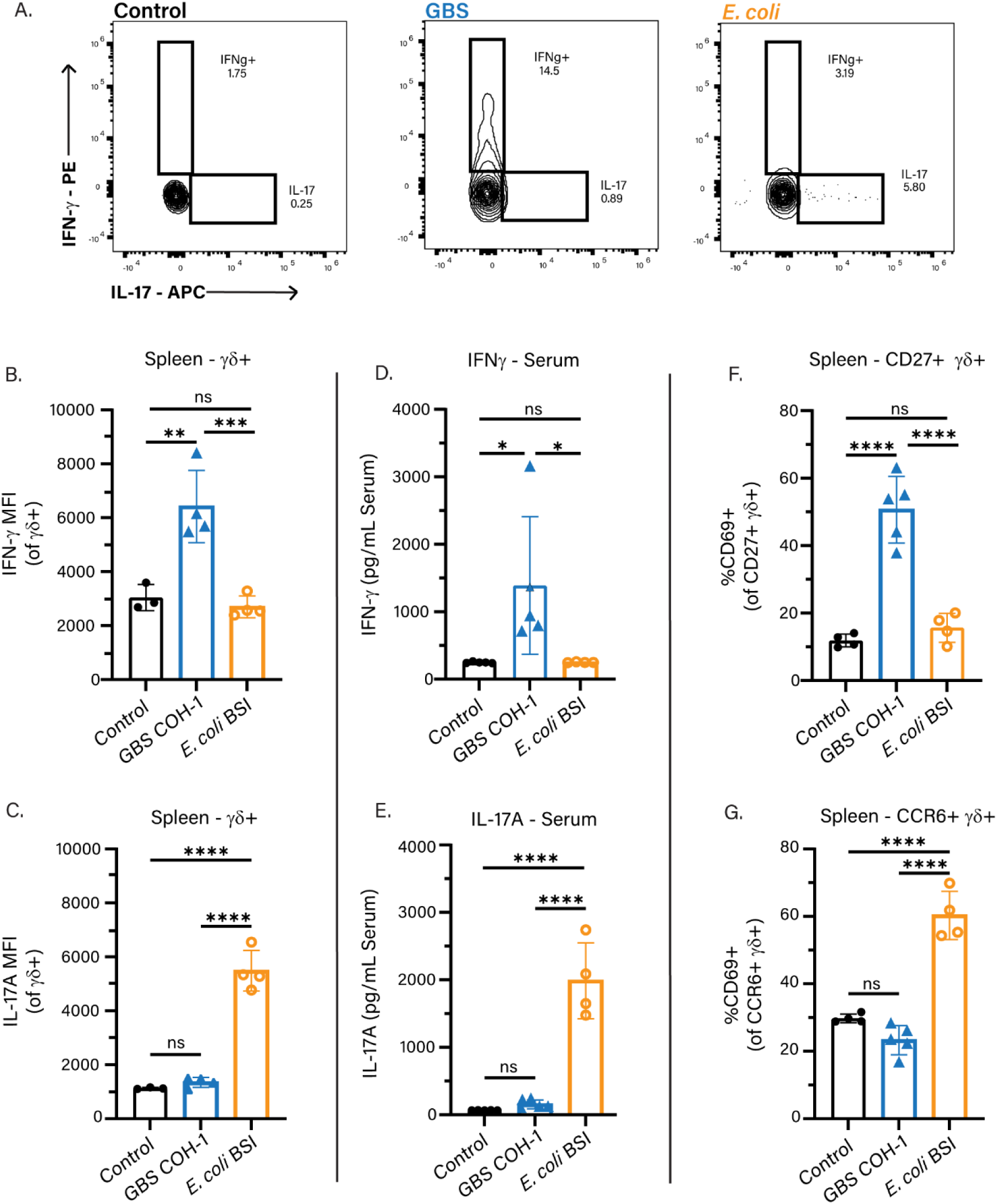
*E. coli* and GBS Neonatal Sepsis Drive Distinct Effector Cytokine Responses from γδ T cells. BL/6 pups were infected with GBS or *E. coli* on P7, and the splenic and systemic cytokine response was measured 18 hours post-infection. A) Flow cytometry gating scheme of IFN-y+ and IL-17+ splenic γδ+ T cells 18 hours post-infection. B) Mean fluorescent intensity (MFI) of IFN-γ and C) IL-17 from total splenic γδ+ T cells 18 hours post-infection. D) IFN-γ and E) IL-17 serum ELISA 18 hours post-infection, samples were pooled from infected pups from A and B. F) Proportion of activated CD27+ and G) CCR6+ γδ+ T cells from uninfected, GBS and *E. coli* infected BL/6 P7 pups 18 hours post-infection. Data shown is from three independent experiments, n>3 mice per group, where each dot represents one mouse. Uninfected age-matched littermates were used as controls. Statistical tests used include one-way ANOVA (B-G) with ns = p>0.05, * = p ≤ 0.05, ** = p ≤ 0.01, *** = p ≤ 0.001, **** = p ≤ 0.0001.

### E. coli *Neonatal Infection Drives Pathogenic γδTCR Signaling*

γδ T cells are capable of undergoing activation via multiple pathways, including MHC-independent TCR activation by pathogen-derived non-peptide antigens.^30^ Nur77 is a transcription factor that is rapidly and specifically expressed during antigen-receptor-mediated signaling and activation in T and B cells.^31,32^ Therefore, we utilized P7 neonatal Nur77-GFP reporter pups to determine if γδTCR signaling occurs during GBS and *E. coli* sepsis. In the spleens of *E. coli*-infected pups 18 hours post-infection, there was an increase in the proportion of Nur77+ CD69+ γδ T cells (Fig. 3a, b), indicating γδTCR-mediated activation. Importantly, nearly all Nur77+ CD69+ γδ T cells expressed CCR6 (Fig. 3c), suggesting that the IL-17 signature during *E. coli* neonatal sepsis is associated with γδTCR signaling. To gain insight into the nature of the *E. coli* antigen recognized by γδ T cells, we performed an *in vitro* culture of heat-killed (HK) *E. coli* BSI-B with Nur77-GFP splenocytes. Culture with heat-killed *E. coli* BSI-B was sufficient to increase the proportion of Nur77+ CD69+ γδ T cells (Fig. S2a, b), suggesting that live *E. coli* is not required for neonatal γδTCR activation. We next sought to determine if γδTCR signaling during *E. coli* infection impacts IL-17 production and survival. To this end, we treated pups with 15 μg/g anti-TCRγδ UC7-13D5 antibody (Fig. S3a), which has been shown to induce internalization of the γδTCR *in vivo*.^33^ The use of a second anti-γδTCR antibody confirmed the γδTCR was not expressed in the spleen or liver (Fig. S3b). Importantly, treatment with this antibody during *E. coli* infection was sufficient to decrease the production of IL-17 (Fig. 3d), confirming that IL-17 production during *E. coli* infection is driven by γδTCR signaling. Blockade of the γδTCR in *E. coli*-infected pups was sufficient to rescue mortality independent of bacterial burden (Fig. 3e, f), however, TCRδ-/- pups, which are devoid of a γδ T cell population, rapidly succumb to *E. coli* infection, despite similar bacterial burden to isotype and anti-TCRγδ-treated pups (Fig 3e, f).

**Figure 3:**
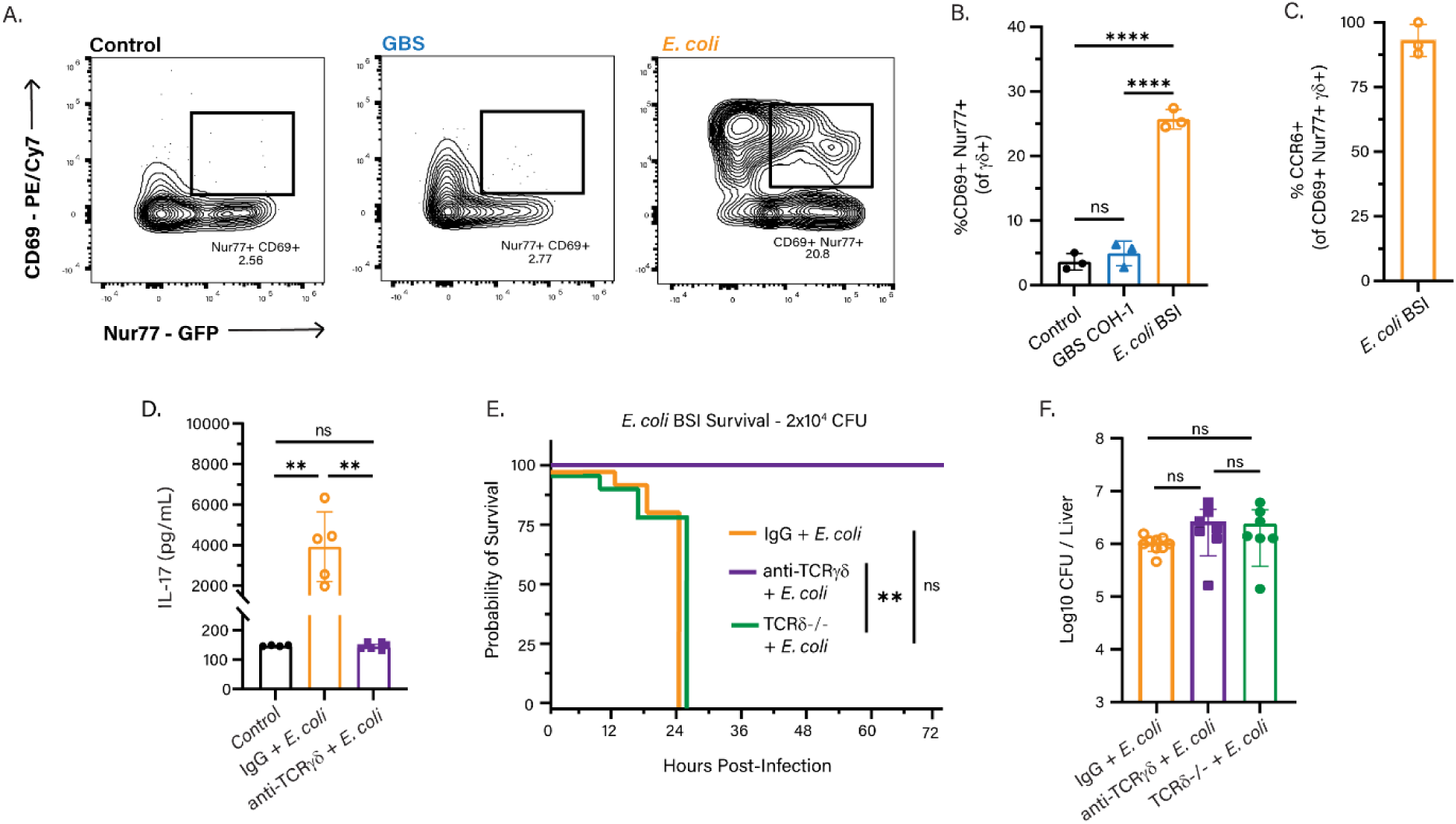
γδTCR Signaling Occurs During *E. coli*, but not GBS Neonatal Sepsis. Nur77^GFP^ pups were infected with GBS or *E. coli* on P7 and spleens were harvested 18 hours post-infection. A) Flow cytometry gating scheme, and B) Quantification of CD69+ Nur77+ splenic γδ+ T cells from control, GBS-infected, and *E. coli-*infected pups. C) Percentage of CD69+ Nur77+ splenic γδ+ T cells from *E. coli* infected pups that express CCR6. D) IL-17 serum ELISA data from *E. coli*-infected pups treated with either isotype IgG or 15 µg/g anti-TCRγδ UC7-13D5 antibody. E) Survival curves and F) Bacterial CFUs from the livers of *E. coli*-infected TCRδ-/- or BL/6 P7 pups treated with either isotype IgG or 15 µg/g anti-TCRγδ UC7-13D5 antibody. Controls are uninfected age-matched littermates. Data shown is from three independent experiments, n>3 mice per group, where each dot represents one mouse. Statistical tests used include Kaplan-Meier (E), one-way ANOVA (B, D, F) with ns = p>0.05, ** = p ≤ 0.01, **** = p ≤ 0.0001.

Interestingly, there was no change in the proportion of Nur77+ CD69+ γδ T cells in the spleens of GBS-infected pups (Fig. 3a, b), suggesting TCR-independent activation of CD27+ γδ T cells during GBS infection. As such, both blockade of the γδTCR, and global deletion of γδ T cells had no impact on survival or pathogen burden during GBS infection (Fig. S3c, d). Therefore, we next asked if IFN-γ production from γδ T cells during GBS infection was driven by other cytokines in the inflammatory milieu. γδ T cells can produce IFN-γ in response to IL-12 and IL-18 ^29,34^; therefore, we tested the serum of GBS-infected pups for the presence of these cytokines. During neonatal GBS infection, no statistically significant increases in IL-12 or IL-18 were observed (Fig. S4a-c), therefore, it is unlikely that IFN-γ production during GBS infection is driven by these cytokines. Overall, these data suggest that the CCR6+ IL-17+ γδ T cells responding to *E. coli* and CD27+ IFN-γ+ γδ T cells responding to GBS neonatal sepsis undergo distinct pathways of activation to elicit effector cytokine production.

### Neuroinflammation is a Feature of E. coli and GBS Neonatal Sepsis

Adverse neurologic outcomes are associated with inflammatory events in early life, including bacterial sepsis.^9,35^ We therefore sought to characterize the neuroinflammatory phenotypes associated with *E. coli* and GBS septicemia. Live *E. coli* and GBS were present in the brains of P7 pups 18 hours post-infection (Fig. 4a), suggesting neuroinflammation during neonatal sepsis is not the result of sterile inflammation, but is rather driven by live bacteria in the brain parenchyma. We next sought to characterize changes to immune cell populations in the neonatal brain during GBS and *E. coli* infection. Flow cytometry analysis of CD45^hi^ immune cells in the perfused brains of P7 pups revealed a significant increase in monocytes and neutrophils in the brains of *E. coli*-infected pups compared to uninfected control pups and GBS-infected pups (Fig. 4b, c, Fig. S5). Interestingly, no significant increase in brain-infiltrating monocytes or neutrophils was noted in GBS-infected pups compared to control mice (Fig. 4b, c).

**Figure 4:**
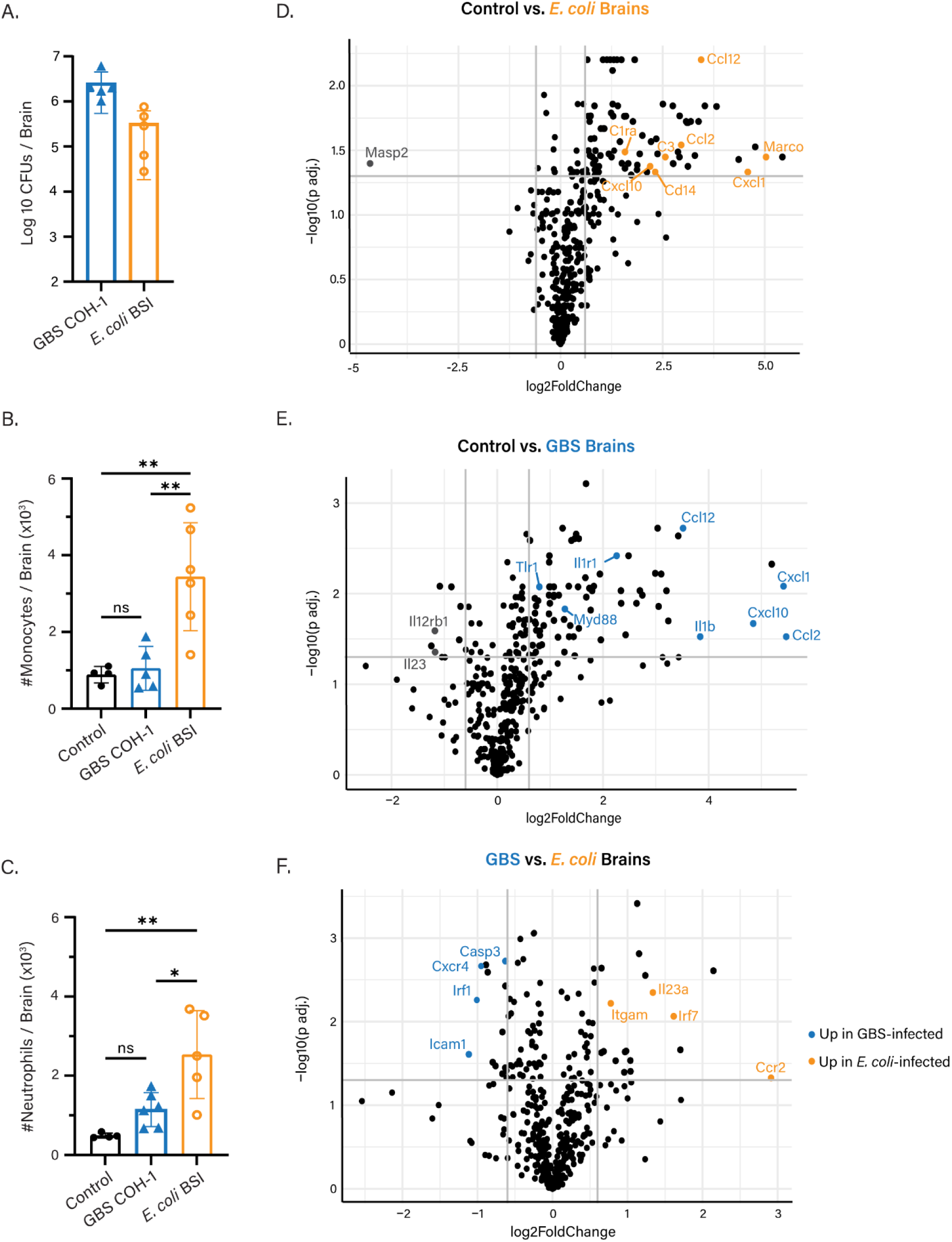
Neuroinflammation is a Feature of *E. coli* and GBS Neonatal Sepsis. BL/6 P7 pups were infected with GBS or *E. coli* on P7. A) GBS and *E. coli* CFUs from the perfused brains of *E. coli*-infected or GBS-infected pups 18 hours post-infection. B) Absolute number of monocytes (CD45^hi^, CD11b+, Ly6C+, Ly6G^low^) and C) Neutrophils (CD45^hi^, CD11b+, Ly6C^low^, Ly6G+) in the perfused brains of *E. coli*-infected or GBS-infected 18 hours post-infection. D) Volcano plot of differentially expressed genes between BL/6 uninfected control vs. *E. coli* infected pups and E) Volcano plot of differentially expressed genes between BL/6 uninfected control vs. GBS infected P7 pups. Data shown is from four independent experiments, n>4 mice per group. Controls are uninfected age-matched littermates. Statistical tests used include one-way ANOVA (B, C) with ns = p>0.05, * = p ≤ 0.05, ** = p ≤ 0.01.

To further investigate the neuroinflammatory phenotype associated with GBS and *E. coli* neonatal sepsis, we measured the mRNA of immunological genes from bulk brain tissue using the Nanostring nCounter Gene Expression platform (Table S1). In the brains of both GBS and *E. coli*-infected P7 pups, there was a significant increase in the expression of monocyte and neutrophil chemotactic factors, such as *Ccl2* and *Cxcl1*, respectively (Fig. 4d, e). Compared to the brains of uninfected control mice, *E. coli*-infected pups had increased expression of genes associated with TLR4, such as *Cd14*, and the complement pathway, such as *C1ra, C3* (Fig. 4d). Compared to uninfected pups, the brains of GBS-infected pups had increased expression of genes involved in the innate immune response to Gram-positive bacteria, such as *Tlr1* and *MyD88* (Fig. 4e). There were thirty-two differentially expressed genes between the brains of GBS and *E. coli*-infected pups including increased expression of *Casp-3* during GBS infection, and increased *Il23a* expression during *E. coli* infection (Fig. 4f). Furthermore, principal component analysis (PCA) of Nanostring data revealed that *E. coli* and GBS-infected brains cluster distinctly from one another (Fig. S6a). Overall, these findings indicate that although both GBS and *E. coli* can cause neuroinflammation during single-organism neonatal sepsis infection, they induce distinct inflammatory phenotypes in the neonatal brain.

### γδ T cells Differentially Impact GBS and E. coli Sepsis-Associated Neuroinflammation

Central nervous system (CNS)-resident γδ T cells are highly skewed toward IL-17 production and CCR6 expression at baseline^36,37^ and play indispensable roles during homeostasis.^36,38^ Under neuroinflammatory conditions, γδ T cells can have both protective ^39^ and pathogenic ^40,41^ effects on the CNS. Therefore, we sought to understand the contribution of γδ T cells to the neuroinflammatory phenotypes observed during GBS and *E. coli* neonatal sepsis. Similar to the spleen, we observed an increase in the proportion of Nur77+ CD69+ γδ T cells in the brain of *E. coli*, but not GBS-infected pups (Fig. 5a). Importantly, the majority of these Nur77+ CD69+ γδ T cells also expressed CCR6 (Fig. 5b), demonstrating that in the brain, IL-17 producing γδ T cells undergo TCR-mediated activation during neonatal *E. coli* sepsis.

**Figure 5:**
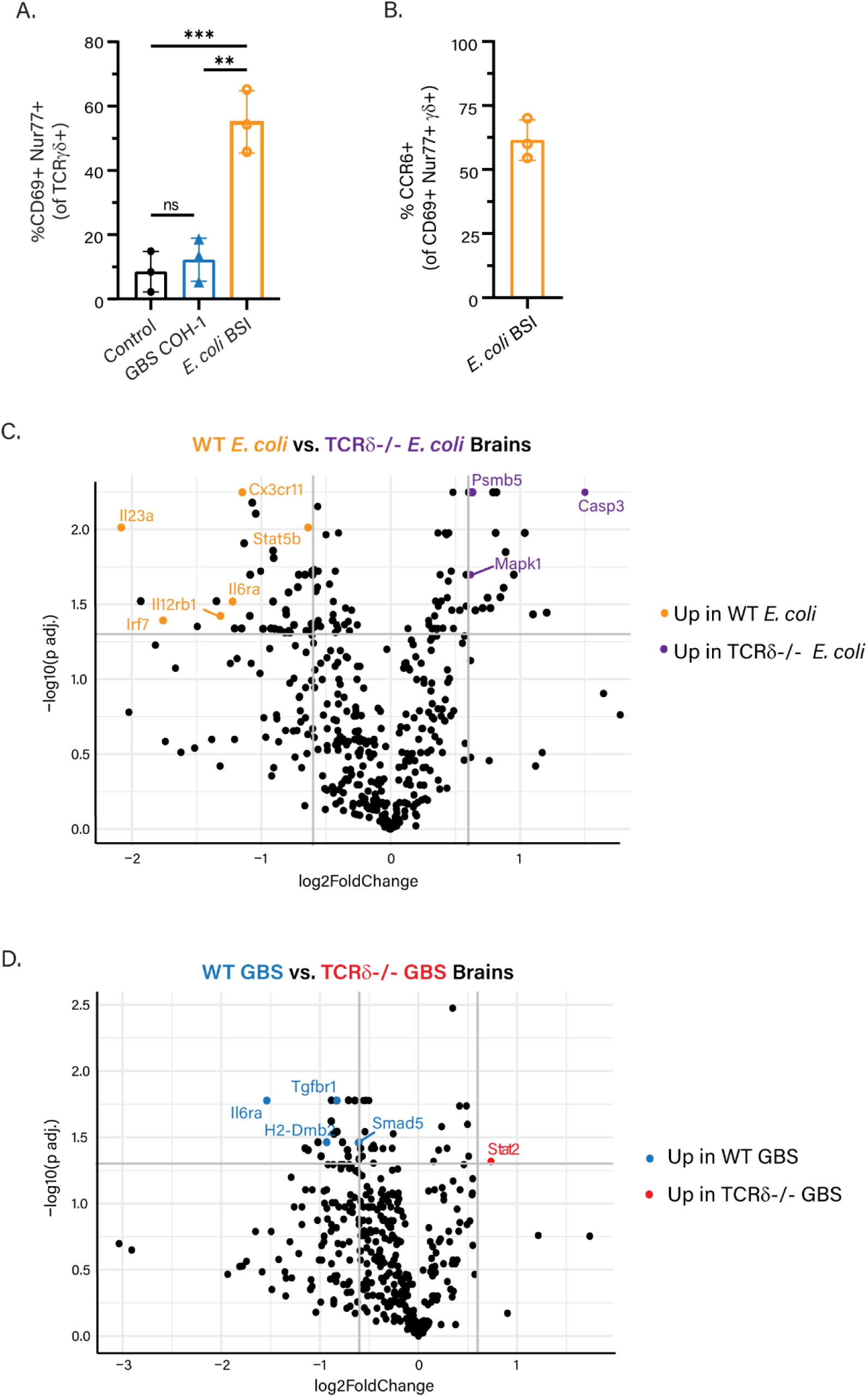
γδ+ T cells Differentially Impact GBS and *E. coli*-Driven Neuroinflammation. Nur77^GFP^ pups were infected with GBS, or *E. coli* on P7, and brains were harvested 18 hours post-infection. A) Proportion of Nur77+ CD69+ γδ+ T cells in the perfused brain of GBS-infected or *E. coli-*infected Nur77-GFP P7 pups. B) CCR6 expression of Nur77+ CD69+ γδ+ T cells in the spleen and brain of *E. coli*-infected pups. (C-D) TCRδ-/- pups were infected with GBS or *E. coli* on P7, and brains were harvested 18 hours post-infection. Volcano plots of differentially expressed genes in the brains of C) *E. coli* infected, or D) GBS-infected BL/6 vs. TCRδ-/- pups. Controls are uninfected age-matched littermates. Data shown is from three independent experiments, n>3 mice per group. Statistics used include one-way ANOVA (A) with ns = p>0.05, ** = p ≤ 0.01, *** = p ≤ 0.001.

To determine the contribution of γδ T cells to GBS and *E. coli* sepsis-associated neuroinflammation, we compared immunological gene expression between BL/6 and TCRδ-/- pups. TCRδ-/- pups were used to evaluate the contribution of γδ T cells to sepsis-induced neuroinflammation, as treatment with the anti-TCRγδ UC7-13D5 antibody did not affect brain-resident γδ T cells (Fig S3b). While there were significant differences in gene expression between uninfected BL/6 and TCRδ-/- pups (Fig. S6b), these conditions clustered similarly to one another, and distinctly from the infection points (Fig. S6a). Analysis of the brain by Nanostring revealed fifty-eight differentially expressed genes between BL/6 and TCRδ-/- *E. coli*-infected pups. Compared to BL/6 *E. coli*-infected pups, TCRδ-/- *E. coli*-infected pups had increased expression of genes associated with apoptosis, such as *Casp-3, Psmb5* and *Mapk1* (Fig. 5c). Notably, uninfected TCRδ-/- pups also had increased *Casp-3* expression compared to uninfected WT BL/6 mice (Fig. S6b), suggesting that γδ T cells may protect the neonatal brain from apoptosis both at baseline and during systemic inflammation. Compared to TCRδ-/- *E. coli*-infected pups, WT BL/6 *E. coli*-infected pups had increased expression of *Il23a*, *Il12rb1*, and *Stat5b* (Fig. 5c). *Il23a and Il12rb1* were also increased in uninfected WT BL/6 pups compared to TCRδ-/- pups, suggesting a role for γδ T cells in driving IL12/IL23 cytokine signaling at baseline and during *E. coli*-mediated systemic inflammation.

During GBS infection, there were twenty-two differentially expressed genes between BL/6 and TCRδ-/- GBS-infected pups. Compared to TCRδ-/- GBS-infected pups, BL/6 pups had increased expression of *H2-Dmb2,* along with *Tgfbr1, Il6ra,* and *Smad5* (Fig. 5d), suggesting a role for γδ T cells in antigen presentation and TGF-β signaling, respectively. Moreover, principal component analysis (PCA) also revealed distinct clustering of TCRδ-/- *E. coli*-infected pups and TCRδ-/- GBS-infected pups (Fig. S6a). Taken together, these data suggest that γδ T cells play an infection-specific role during sepsis-associated neuroinflammation.

## Discussion

The present study reveals that γδ T cells undergo rapid activation and cytokine production during a murine model of single-organism *Streptococcus agalactiae* (GBS) and *Escherichia coli* neonatal sepsis (Fig. 1), and that GBS and *E. coli* septicemia induce activation of IFN-γ and IL-17 producing γδ T cells, respectively (Fig. 2). Notably, a greater proportion and absolute number of γδ T cells were activated in response to *E. coli* infection compared to GBS infection (Fig. 1a-c), despite higher pathogen burden during GBS infection (Fig. 3f, S3c). These findings may suggest that bacterial components, rather than bacterial burden alone, are critical in driving the magnitude of the neonatal γδ T cell response. Therefore, how components of *E. coli* and GBS drive differential γδ T cell activation will be the subject of continued investigation.

Although the γδ T cell compartment had the highest proportion of cells undergoing activation in response to both *E. coli* and GBS infections, a modest increase in activation status was observed for CD4+ and CD8+ T cells, and B cells (Fig. S1a-c). This small proportion of activated CD8+ T cells may represent “virtual memory” (T_VM_) T cells, which are abundant in neonates.^42,43^ Increased activation status of CD8+ T cells at the early timepoint of 18 hours post-infection may further implicate T_VM_ cells, as they are capable of responding to infection faster than naïve T cells due to their memory-like capacity.^44,45^

An additional outstanding question raised by the present findings is the extent to which γδ T cell effector cytokine responses depend directly on bacterial factors such as virulence factors vs. the upstream immune response to such bacterial components. The clinical isolate of *E. coli* used herein is an extraintestinal pathogenic *E. coli* (ExPEC) that expresses several virulence genes, including the capsular polysaccharide K1.^3,7^ The K1 antigen has been implicated in various neonatal infections, including meningitis and sepsis,^46^ and plays a critical role in the ability of *E. coli* to resist phagocytosis and invade the central nervous system.^47^ Similarly, GBS COH-1 (ATCC) is a highly virulent, encapsulated serotype III clinical isolate that expresses virulence factors involved in immune resistance and neuroinvasion, including the serine protease CspA,^48^ and invasion-associated gene (IagA),^49^ respectively. Thus, the extent to which activation and cytokine production from γδ T cells is dependent on bacterial virulence factors vs. signals from the inflammatory milleu requires further investigation.

We also report that γδ T cells undergo distinct pathways of activation to elicit their effector cytokine responses during *E. coli* and GBS sepsis. During GBS infection, there was no change in the proportion of CD69+ Nur77+ γδ T cells in the spleen or brain (Fig. 3a-b, Fig. 5a), suggesting TCR-independent γδ T cell activation and IFN-γ production. As γδTCR signaling was not implicated during GBS infection, the use of the γδTCR blocking antibody during GBS infection conferred no survival benefit or change in pathogen burden (Fig. S3c-d). Similar to NK cells, γδ T cells can produce IFN-γ in response to IL-12, and IL-18,^34,50^ and upon engagement of NKG2D ligands by host stress molecules.^51^ Analysis of the serum from GBS-infected pups revealed no statistically significant increase in IL-12 or IL-18 production (Fig. S4a-c), suggesting that other host stress factors are driving IFN-γ production. Furthermore, the exact mechanism by which CD27+ γδ T cells become activated and make IFN-γ during GBS sepsis will be the subject of future studies.

Conversely, during *E. coli* sepsis, there was a robust increase in the proportion of Nur77+ CD69+ γδ T cells in both the spleen and brain (Fig. 3b, Fig. 5a), suggesting that γδ T cells undergo γδTCR-mediated activation during *E. coli* sepsis. We also report that splenocyte co-culture with heat-killed (HK) *E. coli* was sufficient to induce Nur77 expression in neonatal γδ T cells (Fig. S2), however, the nature of the antigen driving γδ TCR-mediated activation remains unknown. Thus, whether splenic γδ T cells are directly recognizing a heat-stable *E. coli* bacterial product or a host stress molecule that is upregulated in response to both live and heat-killed bacteria, will be the subject of future studies.

Interestingly, we also find that γδTCR signaling during *E. coli* neonatal sepsis is pathogenic, as blocking the γδTCR with the UC7-13D5 antibody was sufficient to rescue mortality (Fig. 3e, Fig. S3a). γδTCR signaling during *E. coli* infection was further associated with the production of IL-17, as a majority of the TCR-activated γδ T cells in the spleen and brain expressed CCR6 (Fig. 3c, Fig. 5b). Moreover, blockade of the γδTCR during *E. coli* infection was sufficient to ablate the IL-17 serum signature seen in isotype-treated pups (Fig. 3d). Improved survival of *E. coli*-infected pups upon blockade of the γδTCR may be due to the reduced production of IL-17 (Fig. 3d, e), as IL-17 has been shown to drive mortality in other models of neonatal sepsis.^52^ Intriguingly, the complete rescue of mortality upon the blockade of γδ T cell functions contrasts with other early-life infections in which γδ T cells are protective to the neonate, including neonatal influenza ^23^ and *Clostridium difficile* infection ^24^. The conflicting role of γδ T cells as pathogenic during *E. coli* sepsis vs. protective during other neonatal infections could be due to the state of immune dysregulation and hyperinflammation that occurs during sepsis, however, the exact nature of these signals, and their specific impact on γδ T cell function, remain unknown. Additionally, these findings raise important questions surrounding the impact of systemic vs. local, tissue-specific γδ T cell responses, and whether a pathogenic contribution of γδ T cells is more likely during a systemic infection.

γδTCR signaling from IL-17+ γδ T cells in the spleen and brain during *E. coli* sepsis also raises interesting questions surrounding γδ T cell subsets and TCR specificity across different organs. γδ T cells develop in discrete sequential waves based on the usage of γ and δ chain pairings, which determine both their effector function and tissue-homing properties.^18^ The γδTCR repertoire and V(D)J diversity vary across subsets, with CNS-resident IL-17+ γδ T cells primarily expressing the invariant Vγ6 chain, and IL-17+ γδ T cells residing outside of the CNS expressing either the Vγ1, Vγ4, or Vγ6 chain, which display varying levels of V(D)J diversity.^29^ Therefore, although there was an increase in the proportion of Nur77+ CD69+ γδ T cells in both the spleen and brain, which subset of IL-17+ γδ T cell is responding to *E. coli* infection in these tissues, along with the nature of the antigen they recognize, remain unknown. Furthermore, whether the γδ T cells in the CNS and periphery recognize the same antigen despite varying TCR repertoire and diversity across these tissues is an interesting direction for future study. Therefore, future analysis will aim to better define the *E. coli* pathogen or host-derived antigen(s) recognized by the γδTCR, along with the contribution of γδ T cell subsets to the unique inflammatory environment in septic tissues.

Post-infectious neurological sequelae represent a significant co-morbidity in survivors of neonatal sepsis.^35^ The present study therefore sought to characterize the unique neuroinflammatory signatures associated with GBS and *E. coli* systemic infection, and the extent to which these signatures are dependent upon γδ T cells. Significant bacterial burden was found in the perfused brains of both GBS and *E. coli*-infected pups (Fig. 4a), demonstrating the capacity of both GBS COH-1 and *E. coli* BSI-B for neuroinvasion. Interestingly, *E. coli* infection induced greater infiltration of monocytes and neutrophils in the brain (Fig. 4b, c), despite both GBS and *E. coli* showing increased gene expression of potent monocyte and neutrophil chemotactic factors, such as *Ccl2*, *Cxcl1,* and *Cxcl10,* respectively (Fig. 4d, e). Differences in neutrophil and monocyte infiltration into the brain during GBS infection may be due to the increased expression of *Casp-3* in the brains of GBS-infected pups compared to *E. coli-*infected pups (Fig. 4f). Furthermore, increased cell death could contribute to the lack of live cellular infiltrate into the brain during GBS infection, as only live cells are used for quantification (Fig. S5). Thus, live monocytes and neutrophils may be recruited to the neonatal brain during GBS infection but may be undergoing apoptosis due to signals in the inflammatory environment. Similarly, which cell type expresses *Casp-3* during GBS infection, and whether this expression is protective or pathogenic, represents an important direction for future study. Furthermore, during *E. coli* sepsis, and at baseline, there was increased *Casp-3* expression in the brains of TCRδ-/- pups compared to BL/6 infected pups, suggesting that γδ T cells in the brain at homeostasis, and during *E. coli* infection may suppress apoptosis (Fig. 5c, Fig. S6b). Whether this anti-apoptotic effect is due to the direct action of γδ T cell-derived IL-17 was not addressed herein, although IL-17 has been shown to have a pro-survival effect on tumor cells ^53^, and B cells ^54^, however, its exact role in this context remains unknown.

Compared to *E. coli* infection, there were fewer differentially expressed genes between BL/6 and TCRδ-/- pups infected with GBS (Fig. 5c, d), which may suggest a more minor role for γδ T cells during GBS compared to *E. coli* neuroinflammation. In the absence of γδ T cells during neonatal GBS infection, there was decreased expression of genes involved in MHC II antigen presentation, including *H2-Dmb2*, which facilitates the removal of CLIP from MHC II molecules.^55,56^ Similarly, *Il6ra*, *Tgfbr1*, and *Smad3* were also increased in the BL/6 compared to TCRδ-/-GBS infected pups, suggesting that during GBS infection, γδ T cells may impact TGF-β signaling (Fig. 5d). Furthermore, mechanisms by which γδ T cells influence MHCII antigen presentation and TGF-β signaling, and how these pathways impact neurological outcomes in septic neonatal mice, represents an important direction for future study. Moreover, future studies utilizing proteomics or single-cell RNA sequencing may help elucidate differential pathways upregulated by γδ T cells in the periphery and brain during *E. coli* and GBS neonatal sepsis.

These data present evidence that γδ T cell responses during neonatal sepsis rely heavily on the initiating pathogen. The finding that context-specific γδ T cell responses differentially impact mortality may have important clinical implications for the treatment of neonatal sepsis. Overall, this work will help elucidate the pathogen-specific contributions of neonatal γδ T cells to sepsis-induced immunopathology and neuroinflammation.

### Limitations of Study

One significant limitation of the present study is the use of TCRδ-/- pups for survival analysis. During *E. coli* infection, neutralization of the γδTCR was sufficient to rescue mortality, however, this was not true for TCRδ-/- pups, who succumbed to *E. coli*-induced mortality at a similar rate as WT pups (Fig. 3e). As γδTCR signaling is also absent in TCRδ-/- pups, the lack of improved survival in TCRδ-/- pups may suggest a physiological perturbation from a developmental lack of γδ T cells.^18^ Additionally, whether there are TCR-independent contributions from γδ T cells during *E. coli* infection remains unclear. WT and TCRδ-/- mice also succumbed to GBS infection at similar rates (Fig. S3d). Thus, whether γδ T cells are pathogenic during GBS infection, or if the lack of improved survival in TCRδ-/- is due to physiologic changes resulting from a lack of γδ T cells, is not robustly addressed herein. Future studies will utilize inducible deletion of the γδ T cell compartment to ameliorate off-target effects of global γδ T cell deletion.

### RESOURCE AVAILABILITY

Lead Contact: For further information and access to resources please address the lead contact, Kathryn Knoop

Materials and Availability: Bacterial strains generated in this study are available upon request to Kathryn Knoop for *E. coli* BSI-B.

Data and code availability: Nanostring data generated in this study is available at Gene Expression Omnibus (https://www.ncbi.nlm.nih.gov/geo/) Accession GSE253301.

### EXPERIMENTAL MODEL AND STUDY PARTICIPANT DETAILS

#### Mice

C57BL/6, TCRδ-/- and Nur77-GFP mice were purchased from The Jackson Laboratory. All animals were bred following the Institutional Animal Care and Use Committee (IACUC) guidelines at Mayo Clinic. Pups were infected on postnatal day 7 (P7) and all sexes were included in the study. Mice were infected intraperitoneally (i.p.) with either a single-organism culture of 2×10^4^ CFU *E. coli* BSI-B (an ST131 sequence type *E. coli*) (Washington University, St. Louis, MO), or 1×10^6^ CFU GBS COH-1 (ATCC) using an insulin syringe (Cardinal Healthcare). Pups were sacrificed 18 hours later. Anti-TCRγδ antibody (UC7-13D5, Biolegend) was given intraperitoneally for *in vivo* blockade of the TCR on γδ T cells (15 μg/g body weight) as described in Fig. 1d.

### METHOD DETAILS

#### Preparation of Bacteria

Clinical *E. coli* isolates were prepared as described previously ^3^, and GBS COH-1 was obtained from ATCC. Single bacterial colonies of GBS and *E. coli* were taken from a streak plate and placed in a 15 mL conical (Fisher Healthcare) containing 5 mL of LB broth (Fisher Healthcare) and placed into a 37°C incubator overnight. The following day, 10 mL of LB broth was measured into a 50 mL conical tube (Fisher Healthcare), and a sterile dropper was used to place 2-3 drops of overnight bacterial stock into fresh LB broth. Bacteria was shaken at 150 rpm at 37°C until an OD of 0.3 was reached. The bacterial culture was spun down at 10200 rpm for 10 minutes (*E. coli*) or 30 minutes (GBS) and the LB supernatant was discarded. The bacterial pellet was resuspended in 10mL of sterile PBS (Life Technologies) and diluted to proper infection dose. To determine *E. coli* and GBS bacterial burden, the liver was harvested and digested in 1mL of sterile PBS (Life Technologies) and 0.5 g of zirconium beads (0.5mm, Fisher Healthcare) in a safe-lock snap cap tube (Fisher Healthcare) and placed into a tissue homogenizer for 5 minutes. The liver homogenate was then serially diluted in sterile PBS (Life Technologies) to achieve a 1:10^5^ dilution (*E. coli*) or 1:10^6^ dilution (GBS). Bacterial homogenate from *E. coli* and GBS-infected pups was plated on either MacConkey agar (Fisher Healthcare) or Tryptic Soy Agar (DIFCO) plates, respectively. CFUs were counted the following day. For heat-killing of bacteria, 1 mL of overnight bacterial stock was placed into an Eppendorf tube (Fisher Healthcare) and placed on a heat block at 55°C for 90 minutes, shaking at 300 rpm.

#### Enzyme-Linked Immunosorbent Assay (ELISA)

Blood was collected from pups and allowed to clot for 45 minutes at room temperature and spun down at 10,000xg for 2 minutes. The serum was collected and was stored at -20°C until use. ELISAs were performed according to manufacturer’s instructions. Serum samples were diluted 1:5 in ELISA diluent buffer (Biolegend). Analysis of serum IL-12 was performed by EVE Technologies (Calgary, AB).

#### Flow Cytometry

Spleens were harvested from mice and were placed in 1 mL of RPMI 1640 (VWR International LLC), and mechanically homogenized using the frosted end of two glass slides (Fisher Healthcare). Spleens were counted using an automated cell counter (Thermo Fisher Scientific).

Cells were pelleted and resuspended in 500 μL FACS buffer (PBS containing 5% human serum (Sigma-Aldrich), 0.5% BSA (Sigma-Aldrich), 0.1% sodium azide) and allowed to block for 20 minutes at 4°C. Surface master mix was made in FACS buffer and staining was performed for 30 minutes in the dark at 4°C. Following surface staining, samples were washed twice with FACS buffer and acquired on the CyTek Northern Lights Spectral Flow Cytometer (Cytek Biosciences).

#### Liver Digestion for Analysis by Flow Cytometry

Animals were euthanized and the liver was harvested and placed into 2 mL RPMI containing 1mg/mL of DNAse I (Sigma-Aldrich, INC), Collagenase II (Life Technologies) and Collagenase IV (Life Technologies). The mixture was placed into a gentleMACS C Tube (Miltenyi Biotech) and placed onto an OctoMACS Tissue Dissociator (Miltenyi Biotech) for 30 minutes at 37°C. Samples were then quenched with 8 mL of fresh RPMI (VWR International LLC) and added to a new 15 mL conical tube (Fisher Healthcare). Samples were pelleted at 1500 rpm for 5 minutes before being counted and blocked in 1 mL of FACS buffer. Following blocking, samples were stained for surface markers and acquired on CyTek Northern Lights Spectral Flow Cytometer (CyTek Biosciences).

#### Intracellular cytokine staining

Cells were placed in 250 µL RPMI supplemented with 10% FBS, 2mM Glutamine (Gibco), 2mM Pyruvate (BioWhittaker), 50 µg/mL Pen/Strep (BioWhittaker), and 0.55 mM 2-ME (Gibco). 1X protein transport inhibitor (Fisher Healthcare) and 1:1000 phorbol 12-myristate 13-acetate (PMA)/Ionomycin (Cell Activation Cocktail, Biolegend) were added, and samples were placed in an incubator at 37°C for four hours. Samples were spun down and resuspended in FACS buffer to block for 15 minutes. Samples were then stained for surface markers for 30 minutes at 4°C before 100 μL of fixative (Life Technologies) was added to each sample for 30 minutes at room temperature. Samples were then washed once with FACS buffer and once with 1X perm buffer (Biolegend) and spun down at 1500 rpm for 5 minutes. Samples were then resuspended in 100 μL 1X perm buffer and intracellular antibodies and were placed at 4°C overnight. The following day, samples were washed twice with FACS buffer and acquired on the CyTek Northern Lights Spectral Flow Cytometer (CyTek Biosciences).

#### Brain Isolation for Analysis by Flow Cytometry

Mice were anesthetized with 10 ug/g of Ketamine/Xylazine mixture and transcardially perfused with 10 mL of cold, sterile PBS (Life Technologies). Brains were digested as previously described^57^. In brief, brains were isolated and placed into a 50 mL conical tube (Fisher Healthcare) containing 5 mL of RPMI (VWR International LLC). The RPMI containing the brains of the mice were then transferred to a 7 mL glass Tenbroeck Dounce homogenizer (Pyrex) and homogenized until the brain was visibly digested (about 10 plunges). The homogenate was then poured through a 70 μm filter into a new 50 mL conical tube, and 10 more mL of RPMI 1640 was added, along with 1 mL of 10X PBS and 9 mL of Percoll (Sigma-Aldrich INC). The 50 mL conical tubes were then placed into a centrifuge and pelleted at 7840xg for 30 minutes at 4°C. Following the spin, the supernatant was fully aspirated off and samples were washed with 50 mL of fresh RPMI 1640 and spun again at 1500 rpm for 10 minutes. Samples were then blocked in FACS buffer for 15 minutes before surface staining was performed at 4°C for 30 minutes.

#### mRNA Isolation from Brains

Brains were homogenized in 1 mL of sterile PBS (Life Technologies) using a Tenbroeck Dounce homogenizer (Pyrex). 100 μL of brain homogenate was used for mRNA isolation with the Qiagen RNeasy Mini Kit according to manufacturer’s instructions. RNA samples were stored at -80°C until ready for use.

#### Nanostring

Nanostring nCounter Mouse Immunology Max Kit was used following mRNA isolation from the brain. RNA hybridization was performed according to Nanostring manufacturer’s instructions. Samples were incubated for 24 hours at 65°C and then were run on the nCounter Prep Station 5s before being placed on the nCounter Digital Analyzer. Raw and normalized counts were collected from ROSALIND platform. Normalized counts were used for PCA plots, which were plotted using ggplot2 and R version 4.2.3. Log2 fold change and p-values for selected comparisons were calculated by ROSALIND platform. False Discovery Rate was calculated from ROSALIND platform using the Benjamini-Yekutieli calculation for multiple test corrections. Selected comparisons were plotted as volcano plots using ggplot2 and R version 4.2.3. A threshold of > +/-0.6 log2 fold change and p-value of <0.05 was selected for plotting differentially expressed genes.

#### Quantification and Statistical Analysis

Student’s unpaired t-test, One-way ANOVA, and Kaplan-Meier tests were conducted using GraphPad Prism (GraphPad Software, Inc.,). Significance is reported as ns = p > 0.05, * = p ≤ 0.05, ** = p ≤ 0.01, *** = p ≤ 0.001, **** = p ≤ 0.0001.

## Acknowledgments

Work was funded by T32 AI170478 (LTW), and R01 DK134366 (KAK).

*Author Contributions:* LTW and KAK conceived the studies and wrote the manuscript. LTW performed animal experiments, ELISAs, flow cytometry, RNA analysis, and data analysis. KGG assisted with animal husbandry and experiments. All authors reviewed the data and manuscript.

## SUPPLEMENTAL ITEMS

**Table S1: Normalized Expression Counts from Nanostring**

## REFERENCES

1. Bergin, S.P., Thaden, J.T., Ericson, J.E., Cross, H., Messina, J., Clark, R.H., Fowler, V.G., Jr., Benjamin, D.K., Jr., Hornik, C.P., and Smith, P.B. (2015). Neonatal Escherichia coli Bloodstream Infections: Clinical Outcomes and Impact of Initial Antibiotic Therapy. Pediatr Infect Dis J 34, 933–936. 10.1097/inf.0000000000000769.

2. Stoll, B.J., Hansen, N.I., Sánchez, P.J., Faix, R.G., Poindexter, B.B., Van Meurs, K.P., Bizzarro, M.J., Goldberg, R.N., Frantz, I.D., 3rd, Hale, E.C., et al. (2011). Early onset neonatal sepsis: the burden of group B Streptococcal and E. coli disease continues. Pediatrics 127, 817–826. 10.1542/peds.2010-2217.

3. Knoop, K.A., Coughlin, P.E., Floyd, A.N., Ndao, I.M., Hall-Moore, C., Shaikh, N., Gasparrini, A.J., Rusconi, B., Escobedo, M., Good, M., et al. (2020). Maternal activation of the EGFR prevents translocation of gut-residing pathogenic Escherichia coli in a model of late-onset neonatal sepsis. Proceedings of the National Academy of Sciences of the United States of America 117, 7941–7949. 10.1073/pnas.1912022117.

4. Segura-Cervantes, E., Mancilla-Ramírez, J., González-Canudas, J., Alba, E., Santillán-Ballesteros, R., Morales-Barquet, D., Sandoval-Plata, G., and Galindo-Sevilla, N. (2016). Inflammatory Response in Preterm and Very Preterm Newborns with Sepsis. Mediators Inflamm 2016, 6740827. 10.1155/2016/6740827.

5. Shane, A.L., Sánchez, P.J., and Stoll, B.J. (2017). Neonatal sepsis. The Lancet 390, 1770–1780. 10.1016/S0140-6736(17)31002-4.

6. Basu, S. (2015). Neonatal sepsis: the gut connection. Eur J Clin Microbiol Infect Dis 34, 215–222. 10.1007/s10096-014-2232-6.

7. Carl, M.A., Ndao, I.M., Springman, A.C., Manning, S.D., Johnson, J.R., Johnston, B.D., Burnham, C.A., Weinstock, E.S., Weinstock, G.M., Wylie, T.N., et al. (2014). Sepsis from the gut: the enteric habitat of bacteria that cause late-onset neonatal bloodstream infections. Clin Infect Dis 58, 1211–1218. 10.1093/cid/ciu084.

8. Heath, P.T., and Jardine, L.A. (2014). Neonatal infections: group B streptococcus. BMJ Clin Evid 2014.

9. Mynarek, M., Bjellmo, S., Lydersen, S., Afset, J.E., Andersen, G.L., and Vik, T. (2021). Incidence of invasive Group B Streptococcal infection and the risk of infant death and cerebral palsy: a Norwegian Cohort Study. Pediatr Res 89, 1541–1548. 10.1038/s41390-020-1092-2.

10. Tavares, T., Pinho, L., and Bonifácio Andrade, E. (2022). Group B Streptococcal Neonatal Meningitis. Clin Microbiol Rev 35, e0007921. 10.1128/cmr.00079-21.

11. Tsafaras, G.P., Ntontsi, P., and Xanthou, G. (2020). Advantages and Limitations of the Neonatal Immune System. Frontiers in Pediatrics 8. 10.3389/fped.2020.00005.

12. Barrios, C., Brawand, P., Berney, M., Brandt, C., Lambert, P.H., and Siegrist, C.A. (1996). Neonatal and early life immune responses to various forms of vaccine antigens qualitatively differ from adult responses: predominance of a Th2-biased pattern which persists after adult boosting. Eur J Immunol 26, 1489–1496. 10.1002/eji.1830260713.

13. Basha, S., Surendran, N., and Pichichero, M. (2014). Immune responses in neonates. Expert Rev Clin Immunol 10, 1171–1184. 10.1586/1744666x.2014.942288.

14. Li, L., Lee, H.H., Bell, J.J., Gregg, R.K., Ellis, J.S., Gessner, A., and Zaghouani, H. (2004). IL-4 utilizes an alternative receptor to drive apoptosis of Th1 cells and skews neonatal immunity toward Th2. Immunity 20, 429–440. 10.1016/s1074-7613(04)00072-x.

15. Semmes, E.C., Chen, J.-L., Goswami, R., Burt, T.D., Permar, S.R., and Fouda, G.G. (2021). Understanding Early-Life Adaptive Immunity to Guide Interventions for Pediatric Health. Frontiers in Immunology 11. 10.3389/fimmu.2020.595297.

16. Chien, Y.H., Meyer, C., and Bonneville, M. (2014). γδ T cells: first line of defense and beyond. Annu Rev Immunol 32, 121–155. 10.1146/annurev-immunol-032713-120216.

17. Vantourout, P., and Hayday, A. (2013). Six-of-the-best: unique contributions of γδ T cells to immunology. Nature Reviews Immunology 13, 88–100. 10.1038/nri3384.

18. Ribot, J.C., Lopes, N., and Silva-Santos, B. (2021). γδ T cells in tissue physiology and surveillance. Nat Rev Immunol 21, 221–232. 10.1038/s41577-020-00452-4.

19. Havran, W.L., and Allison, J.P. (1988). Developmentally ordered appearance of thymocytes expressing different T-cell antigen receptors. Nature 335, 443–445. 10.1038/335443a0.

20. Parker, M.E., and Ciofani, M. (2020). Regulation of γδ T Cell Effector Diversification in the Thymus. Frontiers in Immunology 11. 10.3389/fimmu.2020.00042.

21. Dimova, T., Brouwer, M., Gosselin, F., Tassignon, J., Leo, O., Donner, C., Marchant, A., and Vermijlen, D. (2015). Effector Vγ9Vδ2 T cells dominate the human fetal γδ T-cell repertoire. Proc Natl Acad Sci U S A 112, E556–565. 10.1073/pnas.1412058112.

22. Gibbons, D.L., Haque, S.F., Silberzahn, T., Hamilton, K., Langford, C., Ellis, P., Carr, R., and Hayday, A.C. (2009). Neonates harbour highly active gammadelta T cells with selective impairments in preterm infants. Eur J Immunol 39, 1794–1806. 10.1002/eji.200939222.

23. Guo, X.J., Dash, P., Crawford, J.C., Allen, E.K., Zamora, A.E., Boyd, D.F., Duan, S., Bajracharya, R., Awad, W.A., Apiwattanakul, N., et al. (2018). Lung γδ T Cells Mediate Protective Responses during Neonatal Influenza Infection that Are Associated with Type 2 Immunity. Immunity 49, 531–544.e536. 10.1016/j.immuni.2018.07.011.

24. Chen, Y.S., Chen, I.B., Pham, G., Shao, T.Y., Bangar, H., Way, S.S., and Haslam, D.B. (2020). IL-17-producing γδ T cells protect against Clostridium difficile infection. J Clin Invest 130, 2377–2390. 10.1172/jci127242.

25. Scasso, S., Laufer, J., Rodriguez, G., Alonso, J.G., and Sosa, C.G. (2015). Vaginal group B streptococcus status during intrapartum antibiotic prophylaxis. Int J Gynaecol Obstet 129, 9–12. 10.1016/j.ijgo.2014.10.018.

26. Yadeta, T.A., Worku, A., Egata, G., Seyoum, B., Marami, D., and Berhane, Y. (2018). Vertical transmission of group B Streptococcus and associated factors among pregnant women: a cross-sectional study, Eastern Ethiopia. Infect Drug Resist 11, 397–404. 10.2147/idr.S150029.

27. Travier, L., Alonso, M., Andronico, A., Hafner, L., Disson, O., Lledo, P.M., Cauchemez, S., and Lecuit, M. (2021). Neonatal susceptibility to meningitis results from the immaturity of epithelial barriers and gut microbiota. Cell Rep 35, 109319. 10.1016/j.celrep.2021.109319.

28. Haas, J.D., González, F.H., Schmitz, S., Chennupati, V., Föhse, L., Kremmer, E., Förster, R., and Prinz, I. (2009). CCR6 and NK1.1 distinguish between IL-17A and IFN-gamma-producing gammadelta effector T cells. Eur J Immunol 39, 3488–3497. 10.1002/eji.200939922.

29. Muñoz-Ruiz, M., Sumaria, N., Pennington, D.J., and Silva-Santos, B. (2017). Thymic Determinants of γδ T Cell Differentiation. Trends Immunol 38, 336–344. 10.1016/j.it.2017.01.007.

30. Constant, P., Davodeau, F., Peyrat, M.A., Poquet, Y., Puzo, G., Bonneville, M., and Fournié, J.J. (1994). Stimulation of human gamma delta T cells by nonpeptidic mycobacterial ligands. Science 264, 267–270. 10.1126/science.8146660.

31. Ashouri, J.F., and Weiss, A. (2017). Endogenous Nur77 Is a Specific Indicator of Antigen Receptor Signaling in Human T and B Cells. J Immunol 198, 657–668. 10.4049/jimmunol.1601301.

32. Moran, A.E., Holzapfel, K.L., Xing, Y., Cunningham, N.R., Maltzman, J.S., Punt, J., and Hogquist, K.A. (2011). T cell receptor signal strength in Treg and iNKT cell development demonstrated by a novel fluorescent reporter mouse. J Exp Med 208, 1279–1289. 10.1084/jem.20110308.

33. Koenecke, C., Chennupati, V., Schmitz, S., Malissen, B., Förster, R., and Prinz, I. (2009). In vivo application of mAb directed against the gammadelta TCR does not deplete but generates “invisible” gammadelta T cells. Eur J Immunol 39, 372–379. 10.1002/eji.200838741.

34. Silva-Santos, B., Mensurado, S., and Coffelt, S.B. (2019). γδ T cells: pleiotropic immune effectors with therapeutic potential in cancer. Nat Rev Cancer 19, 392–404. 10.1038/s41568-019-0153-5.

35. Stoll, B.J., Hansen, N.I., Adams-Chapman, I., Fanaroff, A.A., Hintz, S.R., Vohr, B., Higgins, R.D., Health, N.I.o.C., and Human Development Neonatal Research Network, f.t. (2004). Neurodevelopmental and Growth Impairment Among Extremely Low-Birth-Weight Infants With Neonatal Infection. JAMA 292, 2357–2365. 10.1001/jama.292.19.2357.

36. Ribeiro, M., Brigas, H.C., Temido-Ferreira, M., Pousinha, P.A., Regen, T., Santa, C., Coelho, J.E., Marques-Morgado, I., Valente, C.A., Omenetti, S., et al. (2019). Meningeal γδ T cell-derived IL-17 controls synaptic plasticity and short-term memory. Sci Immunol 4. 10.1126/sciimmunol.aay5199.

37. Wo, J., Zhang, F., Li, Z., Sun, C., Zhang, W., and Sun, G. (2020). The Role of Gamma-Delta T Cells in Diseases of the Central Nervous System. Frontiers in Immunology 11. 10.3389/fimmu.2020.580304.

38. Park, J.H., Kang, I., and Lee, H.K. (2022). γδ T Cells in Brain Homeostasis and Diseases. Front Immunol 13, 886397. 10.3389/fimmu.2022.886397.

39. Gentles, A.J., Newman, A.M., Liu, C.L., Bratman, S.V., Feng, W., Kim, D., Nair, V.S., Xu, Y., Khuong, A., Hoang, C.D., et al. (2015). The prognostic landscape of genes and infiltrating immune cells across human cancers. Nat Med 21, 938–945. 10.1038/nm.3909.

40. Gelderblom, M., Arunachalam, P., and Magnus, T. (2014). γδ T cells as early sensors of tissue damage and mediators of secondary neurodegeneration. Front Cell Neurosci 8, 368. 10.3389/fncel.2014.00368.

41. Welte, T., Lamb, J., Anderson, J.F., Born, W.K., O’Brien, R.L., and Wang, T. (2008). Role of two distinct gammadelta T cell subsets during West Nile virus infection. FEMS Immunol Med Microbiol 53, 275–283. 10.1111/j.1574-695X.2008.00430.x.

42. Akue, A.D., Lee, J.Y., and Jameson, S.C. (2012). Derivation and maintenance of virtual memory CD8 T cells. J Immunol 188, 2516–2523. 10.4049/jimmunol.1102213.

43. Schüler, T., Hämmerling, G.J., and Arnold, B. (2004). Cutting edge: IL-7-dependent homeostatic proliferation of CD8+ T cells in neonatal mice allows the generation of long-lived natural memory T cells. J Immunol 172, 15–19. 10.4049/jimmunol.172.1.15.

44. Haluszczak, C., Akue, A.D., Hamilton, S.E., Johnson, L.D., Pujanauski, L., Teodorovic, L., Jameson, S.C., and Kedl, R.M. (2009). The antigen-specific CD8+ T cell repertoire in unimmunized mice includes memory phenotype cells bearing markers of homeostatic expansion. J Exp Med 206, 435–448. 10.1084/jem.20081829.

45. Lee, J.-Y., Hamilton, S.E., Akue, A.D., Hogquist, K.A., and Jameson, S.C. (2013). Virtual memory CD8 T cells display unique functional properties. Proceedings of the National Academy of Sciences 110, 13498–13503. doi:10.1073/pnas.1307572110.

46. Kaczmarek, A., Budzyńska, A., and Gospodarek, E. (2014). Detection of K1 antigen of Escherichia coli rods isolated from pregnant women and neonates. Folia Microbiol (Praha) 59, 419–422. 10.1007/s12223-014-0315-5.

47. Hoffman, J.A., Wass, C., Stins, M.F., and Kim, K.S. (1999). The capsule supports survival but not traversal of Escherichia coli K1 across the blood-brain barrier. Infect Immun 67, 3566–3570. 10.1128/iai.67.7.3566-3570.1999.

48. Harris, T.O., Shelver, D.W., Bohnsack, J.F., and Rubens, C.E. (2003). A novel streptococcal surface protease promotes virulence, resistance to opsonophagocytosis, and cleavage of human fibrinogen. J Clin Invest 111, 61–70. 10.1172/jci16270.

49. Doran, K.S., Engelson, E.J., Khosravi, A., Maisey, H.C., Fedtke, I., Equils, O., Michelsen, K.S., Arditi, M., Peschel, A., and Nizet, V. (2005). Blood-brain barrier invasion by group B Streptococcus depends upon proper cell-surface anchoring of lipoteichoic acid. J Clin Invest 115, 2499–2507. 10.1172/jci23829.

50. Sedlak, C., Patzl, M., Saalmüller, A., and Gerner, W. (2014). IL-12 and IL-18 induce interferon-γ production and de novo CD2 expression in porcine γδ T cells. Dev Comp Immunol 47, 115–122. 10.1016/j.dci.2014.07.007.

51. Rincon-Orozco, B., Kunzmann, V., Wrobel, P., Kabelitz, D., Steinle, A., and Herrmann, T. (2005). Activation of V gamma 9V delta 2 T cells by NKG2D. J Immunol 175, 2144–2151. 10.4049/jimmunol.175.4.2144.

52. Wynn, J.L., Wilson, C.S., Hawiger, J., Scumpia, P.O., Marshall, A.F., Liu, J.-H., Zharkikh, I., Wong, H.R., Lahni, P., Benjamin, J.T., et al. (2016). Targeting IL-17A attenuates neonatal sepsis mortality induced by IL-18. Proceedings of the National Academy of Sciences 113, E2627.

53. Nam, J.S., Terabe, M., Kang, M.J., Chae, H., Voong, N., Yang, Y.A., Laurence, A., Michalowska, A., Mamura, M., Lonning, S., et al. (2008). Transforming growth factor beta subverts the immune system into directly promoting tumor growth through interleukin-17. Cancer Res 68, 3915–3923. 10.1158/0008-5472.Can-08-0206.

54. Xu, S., and Cao, X. (2010). Interleukin-17 and its expanding biological functions. Cell Mol Immunol 7, 164–174. 10.1038/cmi.2010.21.

55. Doebele, R.C., Busch, R., Scott, H.M., Pashine, A., and Mellins, E.D. (2000). Determination of the HLA-DM interaction site on HLA-DR molecules. Immunity 13, 517–527. 10.1016/s1074-7613(00)00051-0.

56. Santambrogio, L., Berendam, S.J., and Engelhard, V.H. (2019). The Antigen Processing and Presentation Machinery in Lymphatic Endothelial Cells. Front Immunol 10, 1033. 10.3389/fimmu.2019.01033.

57. Cumba Garcia, L.M., Huseby Kelcher, A.M., Malo, C.S., and Johnson, A.J. (2016). Superior isolation of antigen-specific brain infiltrating T cells using manual homogenization technique. J Immunol Methods 439, 23–28. 10.1016/j.jim.2016.09.002.

