## Supplementary Figure 1 for "*Streptococcus agalactiae* and *Escherichia coli* Induce Distinct Effector γδ T Cell Responses During Neonatal Sepsis"

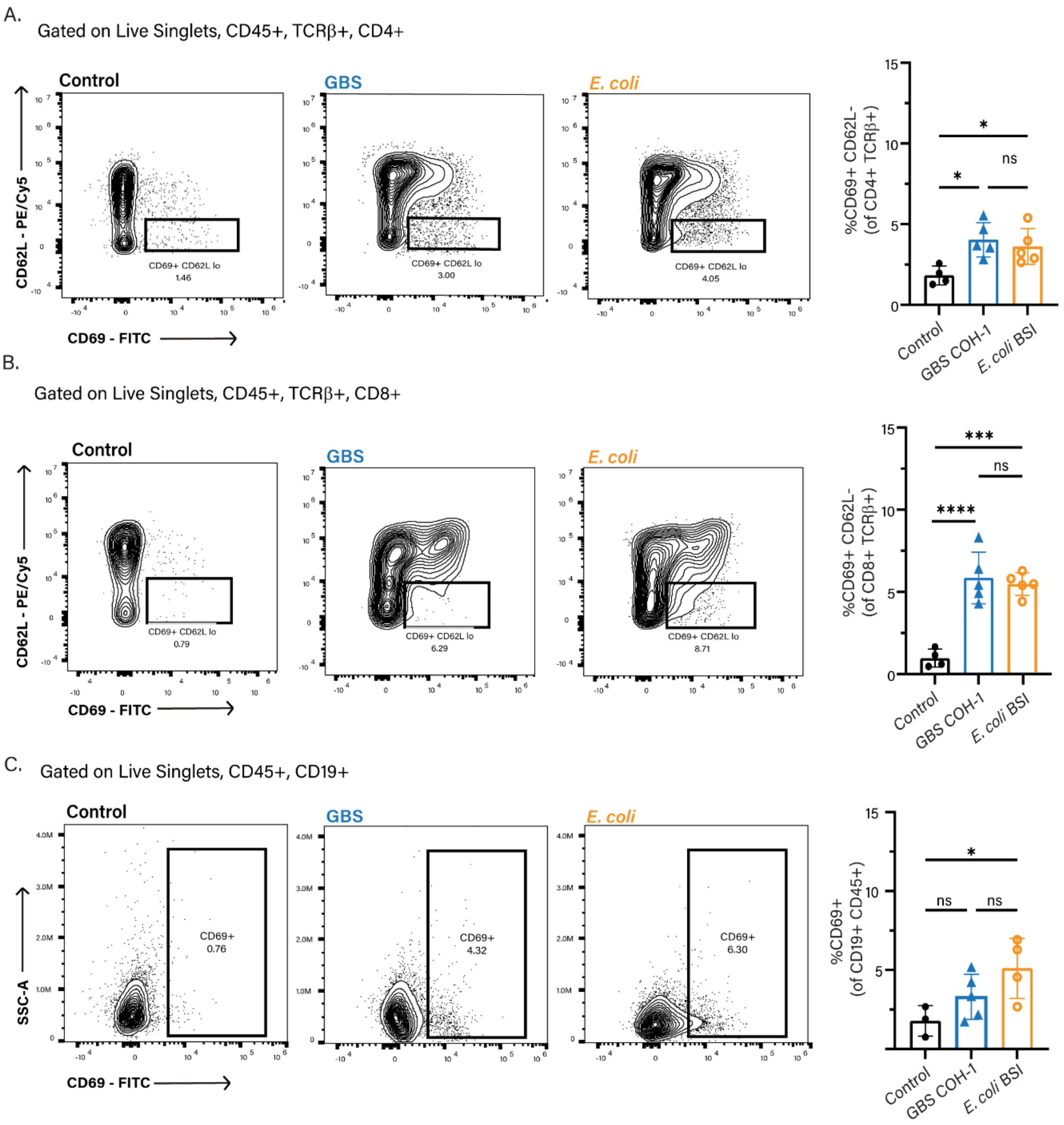


**Figure S1: Responses of CD4+ and CD8+ T Cells, and B cells to GBS and *E. coli* Neonatal Sepsis**

BL/6 pups were infected with GBS or *E. coli* on P7, and the spleen was analyzed 18 hours post-infection. A) Flow cytometric staining of CD69+, CD62L- CD4+ and B) CD8+ T cells and, C) CD69+ B cells. Data shown is from two independent experiments, n>3 mice per group, where each dot represents one mouse. Controls are uninfected age-matched littermates. Statistical tests used include one-way ANOVA (A-C), ns = p>0.05, * = p ≤ 0.05, *** = p ≤ 0.001, **** = p ≤ 0.0001.
