## Supplementary Figure 2 for "*Streptococcus agalactiae* and *Escherichia coli* Induce Distinct Effector γδ T Cell Responses During Neonatal Sepsis"

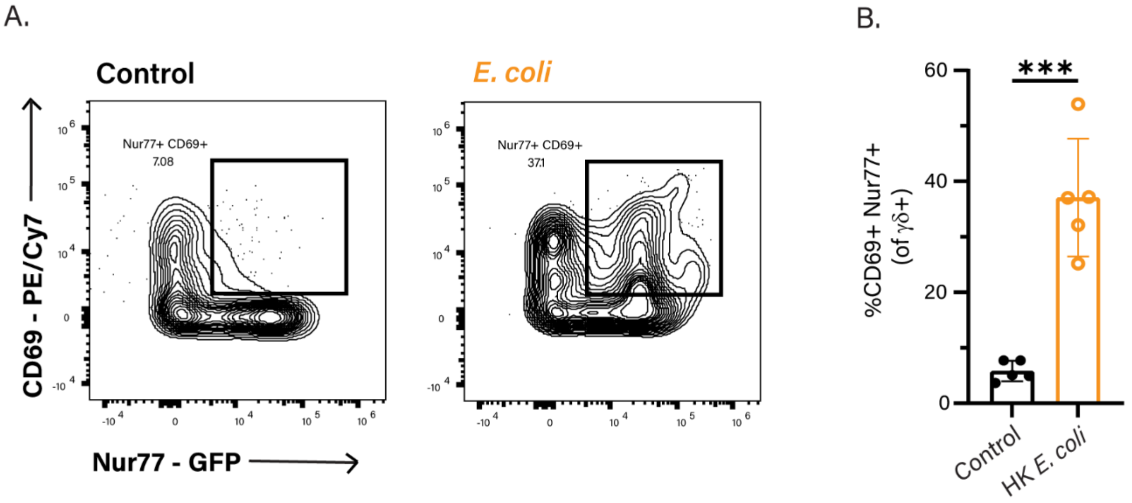


**Figure S2: Heat-Stable E. coli Bacterial Products Lead to γδ TCR Signaling**
Whole splenocytes from P7 Nur77$^{GFP}$ pups were cultured with $10^5$ CFU heat-killed (HK) *E. coli* BSI-B for 24 hours. A) Flow cytometry gating scheme and B) Quantification of CD69+ Nur77+ γδ T cells in control (L) and *E. coli* (R) culture conditions. Data shown is from one experiment, n=5 mice per group, where each dot represents one mouse. Statistical tests used include Student's unpaired t-test (B), with *** = $p \leq 0.001$.
