## Supplementary Figure 3 for "*Streptococcus agalactiae* and *Escherichia coli* Induce Distinct Effector γδ T Cell Responses During Neonatal Sepsis"

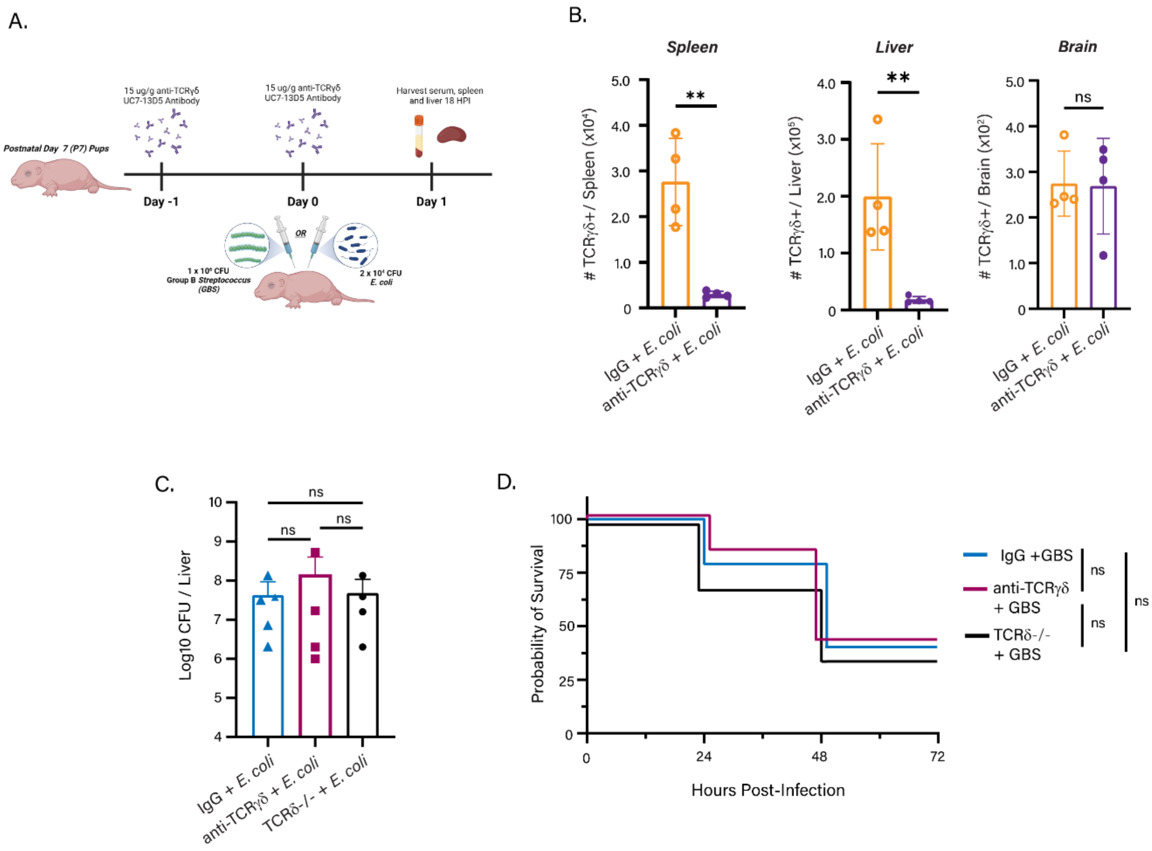


**Figure S3: Effect of Blocking TCRγδ Signaling During GBS Infection**
Mice were treated with either isotype IgG 15 μg/g anti-TCRγδ UC7-13D5 one day before and on the day of infection with either *E. coli* or GBS. A) Experimental schematic B) Staining for TCRγδ in the spleen, liver, and brain upon anti-TCRγδ UC7-13D5 treatment. C) GBS bacterial burden and D) Survival curve of isotype and anti-TCRγδ UC7-13D5 treated C57BL/6 pups, and TCRδ -/-. Controls are uninfected age-matched littermates. Data shown is from three independent experiments, n>3 mice per group, where each dot represents one mouse. Statistical tests used include student's unpaired t test (B), one-way ANOVA (C) Kaplan-Meier (D) with ns = $p>0.05$, ** = $p \leq 0.01$.
