## Supplementary Figure 4 for "*Streptococcus agalactiae* and *Escherichia coli* Induce Distinct Effector γδ T Cell Responses During Neonatal Sepsis"

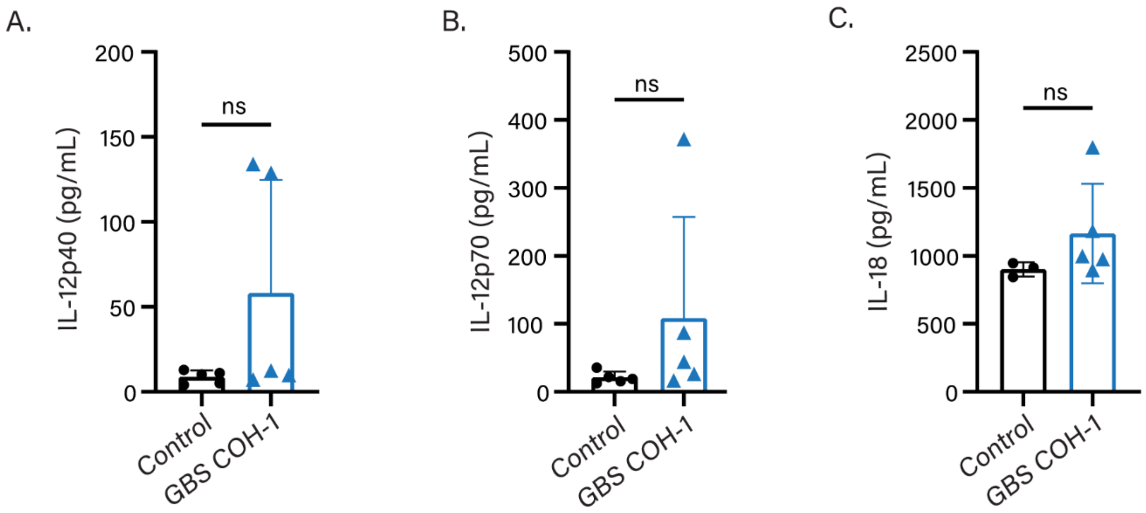


**Figure S4: Serum IL-12 and IL-18 During GBS Neonatal Sepsis**

P7 pups were infected with $10^6$ CFU GBS COH-1 and serum was harvested 18 hours later. A) IL-12p40, B) IL-12p70 serum concentrations obtained from EVE Technologies Multiplex Cytokine Array, and C) IL-18 serum concentrations from ELISA. Data shown is from two independent experiments, n>3 mice per group, where each dot represents one mouse. Controls are uninfected age-matched littermates. Statistical tests used include Student's unpaired t-test (A-C), with ns = $p>0.05$
