## Supplementary Figure 5 for "*Streptococcus agalactiae* and *Escherichia coli* Induce Distinct Effector γδ T Cell Responses During Neonatal Sepsis"

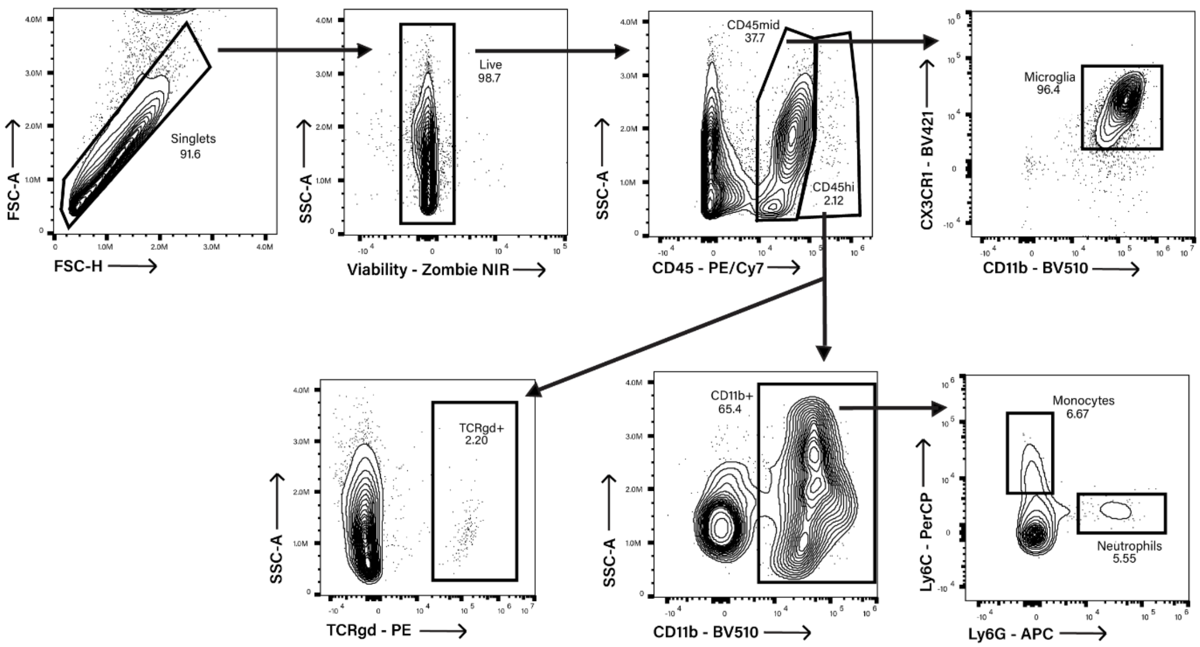


**Figure S5: Flow Cytometry Gating Scheme of Neonatal Brain Immune Cell Populations**

Live, single cells in the brain were gated on CD45, with $CD45^{mid}$ population representing microglia (CD11b+, CX3CR1+). The $CD45^{hi}$ compartment was gated into TCRγδ+ or CD11b+. CD11b+ cells were then gated into Ly6C+, Ly6G- (Monocytes), or $Ly6C^{low}$, Ly6G+ (Neutrophils).
