## Supplementary Figure 6 for "*Streptococcus agalactiae* and *Escherichia coli* Induce Distinct Effector γδ T Cell Responses During Neonatal Sepsis"

A.

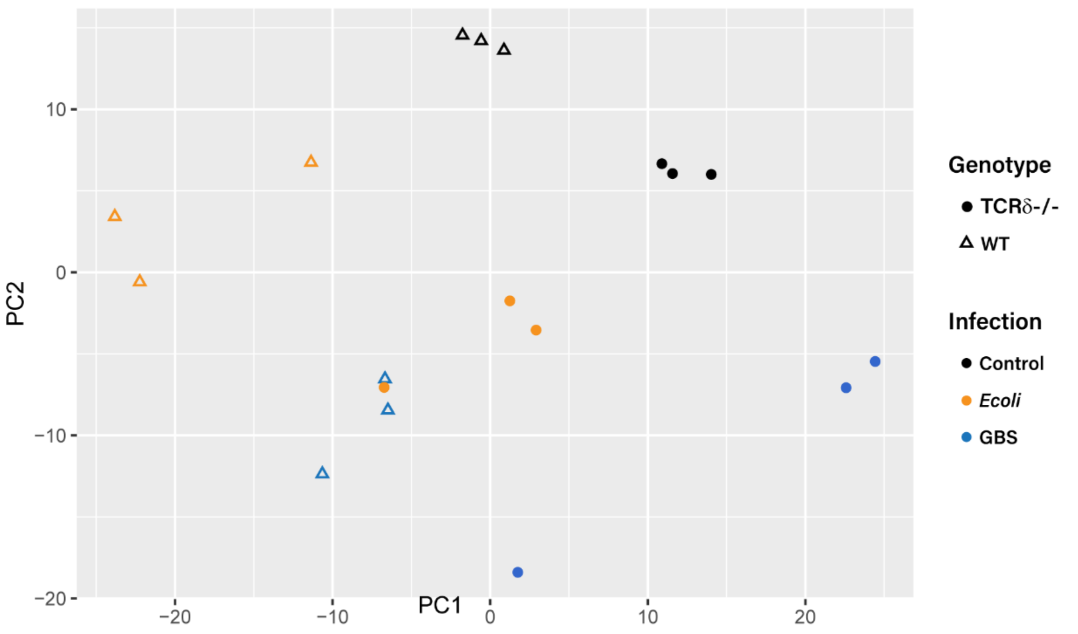


B.

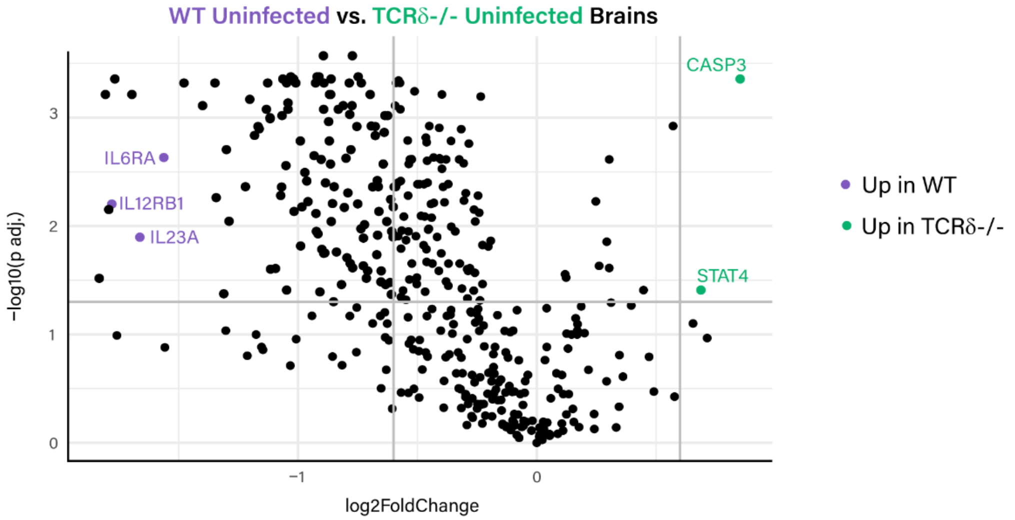


**Figure S6: Principal Component Analysis (PCA) Plot of Gene Expression in Neonatal Brains**

BL/6 pups or TCRδ-/- pups were infected with GBS or *E. coli* on P7, and uninfected age-matched littermates were used as controls. Differentially expressed genes in the brain were assessed by NanoString analysis 18 hours post-infection. A) Principal component analysis was calculated from normalized expression data and clustered

according to genotype (open triangles indicate BL/6 pups or filled circles indicate TCRδ-/- pups) and treatment (uninfected controls, *E. coli* infection, or GBS infection).
B) Volcano plot of differentially expressed genes between WT uninfected controls and TCRδ-/- uninfected controls.
